# Mitochondria regulate *Drosophila* intestinal stem cell differentiation through FOXO

**DOI:** 10.1101/2020.02.12.946194

**Authors:** Fan Zhang, Mehdi Pirooznia, Hong Xu

**Affiliations:** National Heart, Lung, and Blood Institute, National Institutes of Health, USA

**Author notes:** Corresponding author: Hong Xu, Ph.D, National Heart, Lung, and Blood Institute, NIH, Bethesda, 20892 Maryland, USA.

## Abstract

Stem cells often rely on glycolysis for energy production, and switching to oxidative phosphorylation is believed to be essential for their differentiation. To explore the link between mitochondrial respiration and stem cell differentiation, we genetically disrupted electron transport chain (ETC) complexes in the intestinal stem cells (ISCs) of *Drosophila*. We found that ISCs carrying impaired ETC proliferated much more slowly than normal, produced very few intestinal progenitors, or enteroblasts, and failed to differentiate into enterocytes or enteroendocrine cells. One of the main impediments to ISCs’ differentiation appeared to be abnormally elevated forkhead box O (FOXO) signaling in the ETC-deficient ISCs, as genetically suppressing the signaling pathway partially rescued the differentiation defect. Contrary to common belief, neither reactive oxygen species (ROS) accumulation nor adenosine triphosphate (ATP) reduction appeared to mediate the ETC mutant phenotype. Our results demonstrate that ETC is essential for *Drosophila* ISC proliferation and differentiation *in vivo*, and acts at least partially by repressing endogenous FOXO signaling. They also raise the possibility that ETC complexes have a role in stem cell differentiation beyond electron transfer and ATP production.

## INTRODUCTION

Eukaryotes utilize either glycolysis or oxidative phosphorylation (OXPHOS) to extract energy from food and store it as the high-energy ATP molecule. Unlike glycolysis, which partially metabolizes glucose to pyruvate, OXPHOS completely oxidizes glucose to CO_2_, and is much more efficient in harnessing energy (Wallace, 2012). OXPHOS is carried out by the electron transport chain (ETC) complexes and mitochondrial ATP synthase located on the mitochondrial inner membrane. Mitochondria are thought to evolve from endosymbiotic bacteria. Over the course of evolution most mitochondrial genes have migrated to the nucleus, whereas core ETC subunits still remain on the mitochondrial genome (mtDNA) (Wallace, 2012). To accommodate different metabolic profiles and energy demands, cells in different tissues and developmental stages modulate the amount of mtDNA they contain and its expression level (Moraes, 2001). The dynamic regulation of mitochondrial activity is particularly evident during stem cell differentiation. Many types of stem cells favor glycolysis even in the presence of oxygen, a phenomenon termed as “aerobic glycolysis” (Vander Heiden *et al*., 2009; Rafalski *et al*., 2012; Lisowski *et al*., 2018), but undergo a metabolic transition during differentiation (Folmes *et al*., 2012; Zheng, Boyer, *et al*., 2016), increasing mitochondrial mass (Cho *et al*., 2006; Prigione *et al*., 2010; Zhang, Marsboom, *et al*., 2013), mtDNA copy number (Cho *et al*., 2006; Facucho-Oliveira *et al*., 2007; Wanet *et al*., 2014), and respiratory activity (von Heimburg *et al*., 2005; Zhang, Marsboom, *et al*., 2013; Wanet *et al*., 2014). Pharmacological or genetic disruptions of ETC complexes often block stem cell differentiation (Chung *et al*., 2007; Mandal *et al*., 2011), or interfere with lineage specification (Inoue *et al*., 2010; Cherry, Gagne, *et al*., 2013; Diaz-Castro, Pardal, *et al*., 2015), highlighting the importance of ETC activity in stem cell differentiation. However, how ETC complexes might interact with the host cell’s machinery to promote cell differentiation remains unknown.

The *Drosophila* adult midgut is a well-established model to study stem cell behaviors. Intestinal stem cells (ISCs) have been found interspersed among the basal epithelium of the entire adult fly midgut (Buchon, Osman, *et al.*, 2013; Marianes and Spradling, 2013). ISCs proliferate and give rise to new epithelium in response to tissue injury or stresses (Biteau *et al.*, 2011; Jiang and Edgar, 2011; Meng and Biteau, 2015). The ISC usually undergoes an asymmetric division that generates a new ISC and an intermediate progenitor, enteroblast (EB), which can further differentiate into either an absorptive enterocyte (EC) characterized by a polyploid nucleus, or a secretory enteroendocrine (EE) cell at a lower frequency (Ohlstein and Spradling, 2007; Biteau *et al.*, 2008; Jiang and Edgar, 2011; Biteau and Jasper, 2014). ISCs can also undergo symmetric division to proliferate and replenish themselves (de Navascues *et al*., 2012). The simple organization, defined ISC lineage, and ample set of genetic tools available in *Drosophila* make the *Drosophila* midgut an ideal model to explore the impact of ETC complexes on stem cell differentiation.

Applying a selection scheme based on a mitochondrially targeted XhoI endonuclease (MitoXhoI) (Xu *et al*., 2008), we isolated a temperature-sensitive lethal mtDNA mutation (ts), *mt:CoI*^*T300I*^ that abolishes the sole XhoI site in the cytochrome C oxidase (COX) subunit I locus on *Drosophila* mitochondrial genome (Hill, Chen, et al., 2014; Chen et al., 2015). The resulting threonine-to-isoleucine substitution destabilizes the association between the mutated COX1 protein with the cytochrome A heme, which leads to severely reduced cytochrome c oxidase (COX) activity at the restrictive temperature of 29°C (Chen et al., 2015). In tissues or cells harboring homoplasmic ts allele, mitochondrial inner membrane potential is reduced, but no accumulation of reactive oxygen species (Chen *et al*., 2015). Homoplasmic ts flies developed normally at 18°C, but failed to eclose at 29°C. In heteroplasmic flies carrying both the ts mtDNA and wild-type (*wt*) mtDNA, tissue-specific expression of MitoXhoI can effectively remove the wild-type mtDNA, and results in ts homoplasmy in those tissues (Chen *et al*., 2015). In this study, we expressed MitoXhoI in the midgut using the *esg*^*ts*^*Flp-Out (esg*^*ts*^*F/O)* method (Jiang *et al*., 2009). This genetic scheme generated ts homoplasmy in the ISCs and their derivatives in heteroplasmic flies and allowed us to investigate how ETC deficiency impacts ISC differentiation in an overall healthy background. We report that deficiencies in cytochrome chain complexes in midgut stem cells block their proliferation and differentiation, which is partially due to an increase of the activity of the FOXO transcription factor.

## RESULTS

### COX disruption in *Drosophila* ISCs decreases the number of their differentiated progeny cells but not of ISCs

To study how defective respiration might affect the maintenance and differentiation of *Drosophila* ISCs, we generated clones of ISCs harboring the temperature-sensitive lethal mtDNA mutation (*ts*): *mt:CoI*^*T300I*^ (Supplemental Figure S1). In heteroplasmic ts flies, we drove the expression of a mitochondrially targeted XhoI restriction enzyme (*UAS-mitoXhoI*) specifically in ISCs, using the *esg*^*ts*^*F/O* system (*esg-GAL4, tub-GAL80*^*ts*^, *UAS-nlsGFP/CyO; UAS-flp, act>>CD2>>GAL4/TM6*) (Jiang *et al*., 2009), to render the ISCs homoplasmic for the ts mutation (Chen *et al*., 2015). In this system, GAL4 is specifically expressed in ISCs under the esg promoter, but its activity is hampered by ubiquitous GAL80^ts^ at 18°C. Newly emerged adult flies are subjected to a pulse of heat shock at 37°C and maintained at 29°C afterward. This treatment inactivates the GAL80^ts^ protein and allows *esg-Gal4* to activate Flp expression in ISCs (Jiang *et al*., 2009). Flp in turn induces *act>Gal4*, which actives UAS transgenes in all progeny derived from ISCs (Jiang *et al*., 2009). This pulse of heat shock also stresses the midgut to induce ISC proliferation (Strand and Micchelli, 2011). Expression of Mito-XhoI removes the wild-type mtDNA in heteroplasmic cells, rendering them homoplasmic for ts and deficient in COX activity (Chen *et al*., 2015). Meanwhile, defective cells are labeled with GFP and therefore readily distinguished from healthy cells. As a control, we introduced the same combination of nuclear transgenes in a homoplasmic background harboring an synonymous mtDNA mutation that abolishes the XhoI site on *mt:CoI* locus, but does not change the COX amino acid sequence (*mt:CoI*^*syn*^) (Chen *et al*., 2015). From here on out, we refer to the ISCs and their progeny carrying the ts allele as mutant and the ISCs and progeny carrying *mt:CoI*^*syn*^ as control.

In control flies, one third of the midgut was occupied by GFP^+^ cells 3 days after heat shock. The GFP^+^ area increased in older flies, reaching 40% and 60% at day 10 and day 20, respectively (Figure 1, A and B). In the mutants, the midguts were fragile during dissection, and appeared thinner than in controls. Less than 20% of the tissue was GFP^+^ at day 3 (Figure 1B), and the proportion of GFP^+^ area remained the same in older mutant flies (Figure 1B). We stained midguts for Delta, Su(H)GBE-LacZ, Pdm-1 and Prospero to mark ISCs, EB, EC and EE cells, respectively, and determined their abundance in midguts (Supplemental Figure S1, A and C). The abundance of ISCs in the mutant was higher than in control (Supplemental Figure S2A), indicating that COX deficiency does not affect ISCs’ survival and self-renewal. However, the abundance of GFP^+^ differentiated cells (EBs, ECs and EEs) was much less in mutant midguts compared to control (Figure 1, C and D; Supplemental Figure S2A).

**FIGURE 1:**
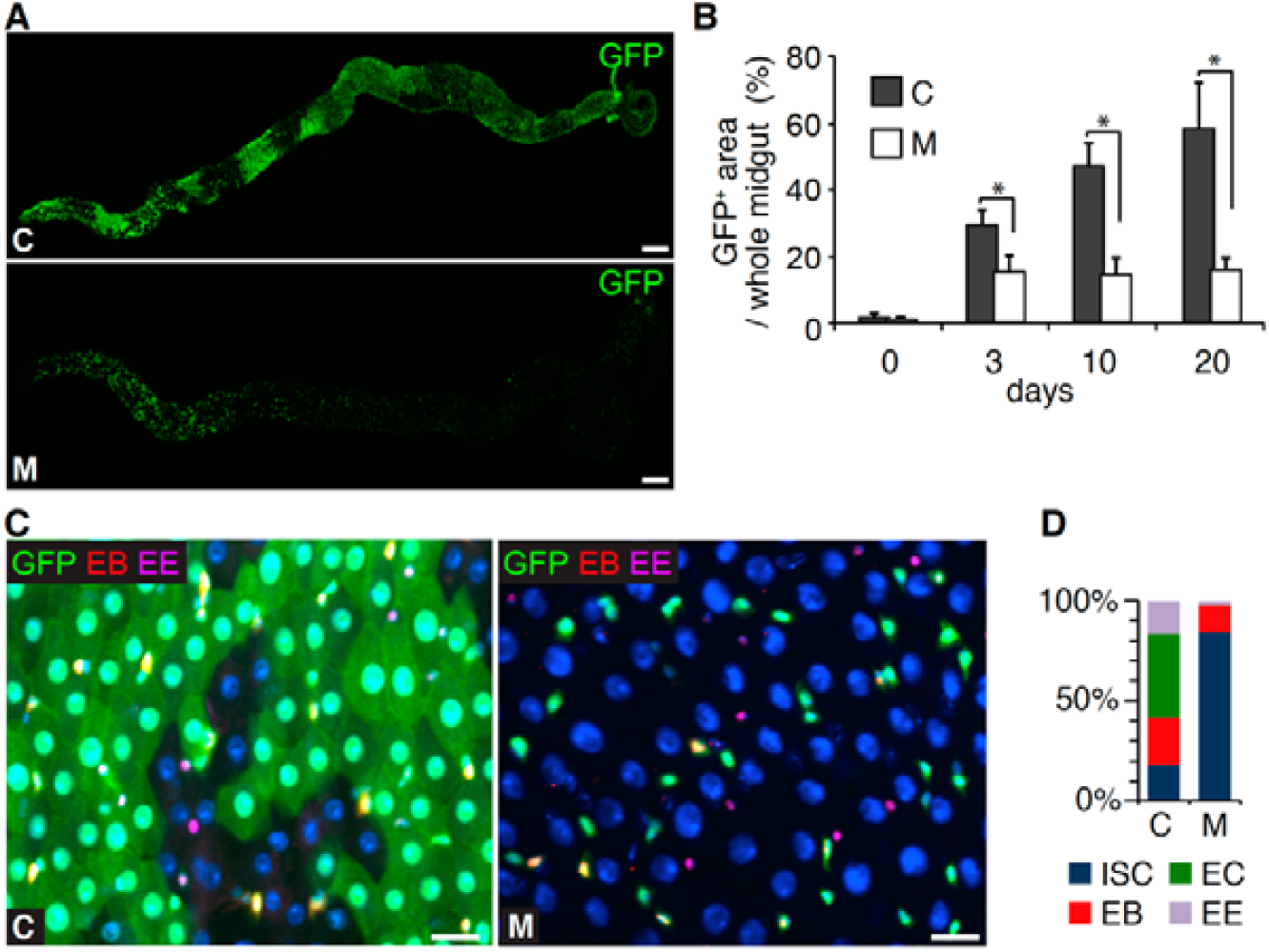
There are less differentiated cells in mutant midgut. (A) Less GFP^+^ cells in the mutant (M) versus control (C) midgut (*esg*^*ts*^*F/O*>*mitoXhoI* flies), 20 days after heat shock. Scale bars: 200 µm. (B) Quantification of the midgut area occupied by GFP^+^ cells in C and M midguts at different time points after heat shock. n=10 midguts each group, error bar: SD. *: p<0.001. (C) Representative views of the C and M posterior midguts, 10 days after the heat shock, showing significantly fewer GFP^+^ EB, EC and EE cells in M than C midgut. Cell-specific markers are: EB: Su(H)GBE-lacZ (red); EC: cell body and nucleus size and EE: Prospero (magenta). Green: GFP. Scale bars: 10 µm. (D) Bar graph showing the composition of GFP^+^ cell types in the C and M midguts, 20 days after the heat shock. n=10 midguts each group, C: 3,545 cells, M: 1,711 cells.

### COX deficiency impairs the proliferation and differentiation of ISCs

Multipotent ISCs proliferate and generate EB cells, which subsequently differentiate into ECs or EE cells. Thus, the paucity of ISC progeny (EB, EC and EE cells) in mutant midguts could be caused by either impaired ISC differentiation; or increased cell death. To test these possibilities, we first assessed apoptosis by staining midguts with an antibody against cleaved-caspase 3, to mark apoptotic cells (Xing *et al*., 2015). Overall, very few cells stained positive for cleaved-caspase 3 in either control or mutant midguts (Figure 2, A and B). They mostly located to the anterior part of midguts and had large nuclei, indicating they were presumably ECs. Of primary significance, the abundance of apoptotic cell was comparable in control and mutant guts (Figure 2, A and B), demonstrating that apoptosis is not increased in mutant midguts, nor should it be considered the cause of the lack of ISC progeny. Additionally, the number of GFP^+^ ISCs were comparable between control and mutant anterior midguts either 8 hours or 3 days after induction (Supplemental Figure S2B), further substantiating that the lack of ISC progeny in the mutant midgut is not caused by an increase of ISC death.

**FIGURE 2:**
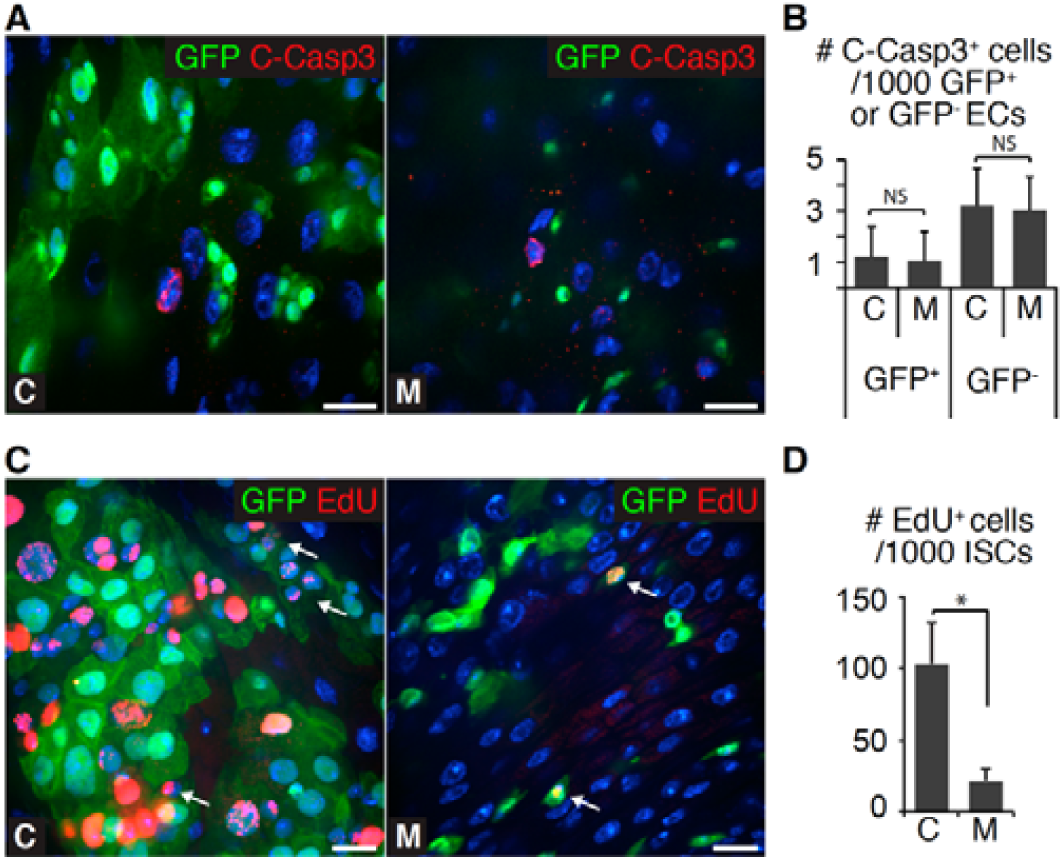
ETC disruption impairs ISC proliferation and EC formation, but does not trigger ISC apoptosis. (A) Cleaved Caspase-3 staining showing equally rare apoptotic cells in both control (C) and mutant (M) midguts (*esg*^*ts*^*F/O*>*mitoXhoI* flies) 20 days after the heat shock. Green: GFP; Red: Cleaved Caspase-3; Blue: DAPI. Scale bars: 10 µm. (B) Quantification of the apoptotic cells marked by Cleaved Caspase-3 staining in C and M midguts, 20 days after the heat shock. n=10 midguts each group; error bar: SD; NS: Not Significant. (C) EdU incorporation (red) in C and M midguts, 20 days after the heat shock. Nuclear EdU incorporation is detectable in GFP^+^ cells with either large or small nucleus in the C midgut, but only in a few small-nucleus GFP^+^ cells in the M midgut. ISC division (arrow) is detectable but at a low frequency in the M midgut. Green: GFP; Red: EdU; Blue: DAPI. Scale bars: 10 µm (D) Quantification of the EdU^+^ ISCs in C and M midguts, 20 days after the heat shock. n=10 midguts each group, 1,000 ISCs assessed per midgut; error bar: SD; *: p<0.001.

To address whether COX deficiency might impair cell proliferation, we examined DNA synthesis *via* EdU incorporation. Many EdU^+^ cells were observed in control midguts (Figure 2C and Supplemental Figure S2C), and the majority of these cells had large nuclei (Figure 2C), indicating they are EB cells undergoing endoreplication to differentiate into EC cells, or newly developed EC cells (Xiang *et al*., 2017). A small fraction of EdU^+^ cells had small nuclei, likely the proliferating ISCs. In mutant midguts, the overall number of EdU^+^ cells were much less than that in controls (Figure 2C and Supplemental Figure S2C). We barely observed EdU^+^ cells with large nuclei (Figure 2C and Supplemental Figure S2C), indicating the lack of EB-to-EC endoreplication in the mutant midgut. Additionally, we only detected very few EdU^+^ cells with small nuclei in mutant midguts (Figure 2, C and D), indicating that the proliferation of mutant ISCs was also impaired. Together, these observations suggest that COX deficiency impairs both ISCs’ proliferation and their differentiation into ECs.

### ETC disruptions block ISC differentiation via pathways independent of mitochondrial energetic deficiency

Active ETC complexes establish mitochondrial inner membrane potential (Δψ_m_) that drives ATP synthesis. To understand the biochemical deficiency in mutant ISCs, we first assessed the Δψ_m_ using ratiometric imaging between TMRM and MitoTracker-deep red, the fluorescent dyes report Δψ_m_ and mitochondrial mass, respectively. Consistent with previous studies (Chen *et al*., 2015; Zhang *et al*., 2019), we found that Δψ_m_, indicated as the ratio of TMRM to Mitotracker was markedly reduced in mutant ISCs compared to the control (Supplemental Figure S3, A and B). Depolarized mitochondria are less effective in producing ATP compared to energized mitochondria with higher Δψ_m_. Indeed, midguts carrying homoplasmic *ts* allele had lower ATP levels compared to control (Supplemental Figure S4A). Noteworthy, ATP deficiency, specifically the reduced ATP-to-AMP ratio activates AMP-activated protein kinase (AMPK), which inhibits cell proliferation through a P53-dependent pathway in a *Drosophila tenured* mutant affecting *coxVa* locus on the nuclear genome (Mandal, Guptan, *et al*., 2005). In addition, RNAi against AMPK can rescue cell proliferation defect in fly eye discs of *tenured* mutant (Mandal, Guptan, *et al*., 2005). However, knocking down AMPK failed to rescue ISCs proliferation defect in mutant midguts (Supplemental Figure S4, B and C), suggesting that a reduced supply of ATP, a common consequence of ETC disruption, is not the main cause of the ISC lineage impairments.

We initially thought that after the elimination of the wild-type mtDNA in heteroplasmic ISCs, the ts mtDNA would undergo compensatory replication to restore the mtDNA content to normal (Supplemental Figure S1B). However, mtDNA content in mutant ISCs remained much lower than that in control cells (Supplemental Figure S5A), suggesting an incomplete compensatory replication of mtDNA. Thus, aside from lacking a functional Complex IV because of the ts mutation, mutant ISCs might also possess inadequate amount of Complexes I, III or V because of an insufficient supply of mtDNA-encoded subunits. To determine whether the differentiation phenotype is specifically pertained to Complex IV, we disrupted each single complex independently by knocking down the nuclear genes encoding their components with RNAi (Figure 3A). We used RNAi lines that were proven effective in a previous study (Teixeira, Sanchez, Hurd, *et al*., 2015), and activated them in ISCs and progeny specifically using the *esg*^*ts*^*F/O* system as described above. RNAis against Complex III and IV nuclear genes caused much stronger defects in ISC differentiation than RNAis against the other complexes (Figure 3B; Supplemental Figure S5, B-D). The greatest impact was observed with disruption of Complex III and was similar to the impact of the mitochondrial *ts* mutant (Figure 3B; Supplemental Figure S5, B-D). Complex III and IV are on the main path of electron transport chain (Figure 3A), while Complex V only uses proton gradient for ATP synthesis and is not directly involved in the electron flow (Figure 3A). Taken together, our results suggest that the blockage of ISC differentiation may not be a simple consequence of ATP deficiency, and instead is likely caused by the defect in electron transport or other specific impairment on Complex III and IV.

**FIGURE 3:**
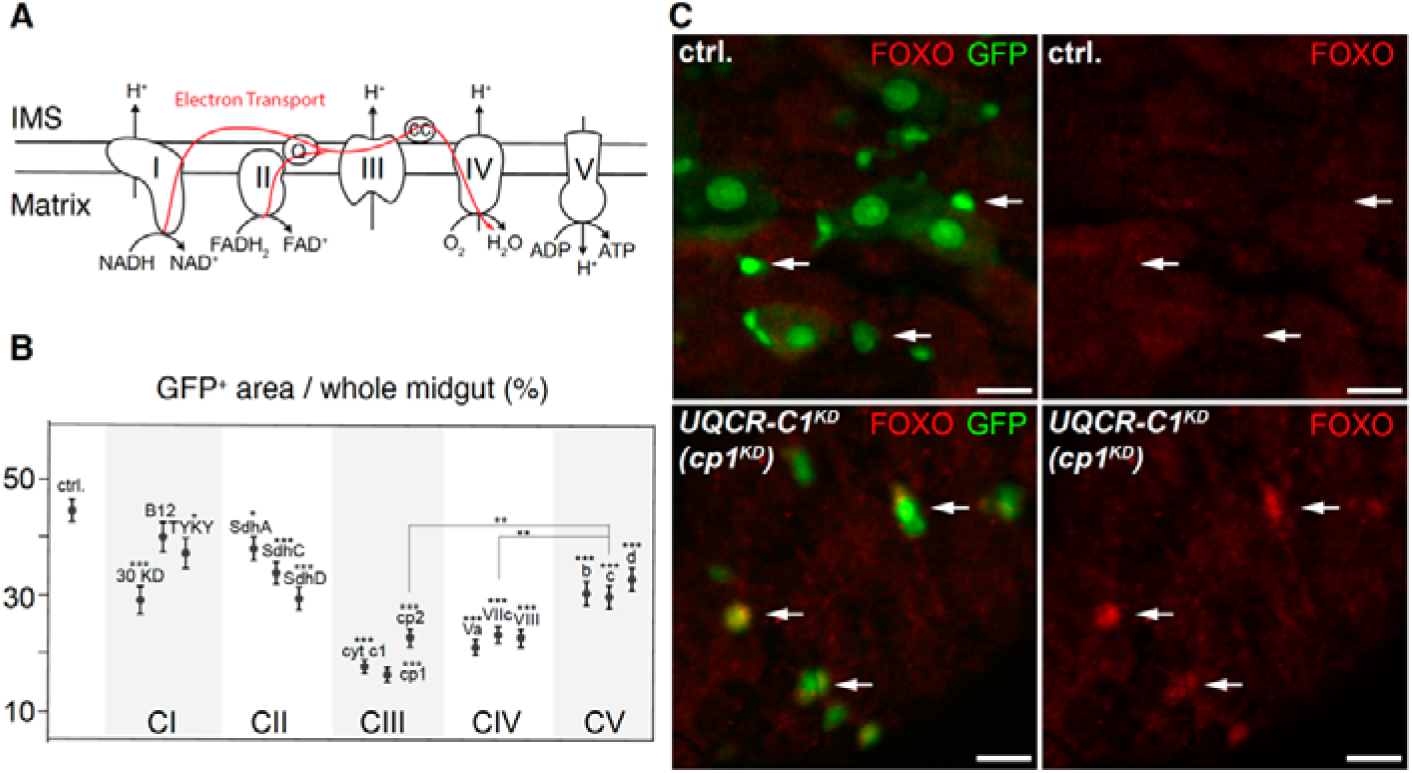
Knockdown of Complex III subunits mimics the *ts* mutant phenotype. (A) Schematic representation of electron transportation and the five mitochondrial respiratory complexes. (B) Quantification of the midgut area occupied by GFP^+^ cells in control and gene knockdown *esg*^*ts*^ *F/O* flies for nuclear-encoded respiratory complex subunits [Complex I: 30KD (ND30), B12(NDB12), TYKY(ND23); Complex II: SdhA (SdhA), SdhC (SdhC), SdhD (SdhD); Complex III: cyt c1(Cytochrome c1), cp1(UQCR-C1), cp2 (UQCR-C2); Complex IV: Va (COX5A), VIIc (COX7C), VIII (COX8); Complex V: b (ATPsynB), c (ATPsynC), d (ATPsynD)], 10 days after heat shock. n=10 midguts each group, error bar: SEM; asterisks above the subunit name indicate the significance compared with RNAi control (*esg*^*ts*^*F/O >luciferase*^*KD*^). *: p<0.05, **: p<0.01, ***: p<0.001. (C) FOXO antibody staining of the control and *UQCR-C1*^*KD*^ *esg*^*ts*^ *F/O* midguts, 10 days after the heat shock. FOXO protein levels dramatically increased in the *UQCR-C1*^*KD*^ ISCs/EBs (arrows) compared with control. Green: GFP; Red: FOXO; Scale bars: 10 µm.

### ETC deficiency blocks the differentiation of ISCs to ECs through the FOXO pathway but independent of ROS

Electron flow from Complex III to Complex IV is a major step generating reactive oxygen species (ROS) (Starkov, 2008; Bleier and Drose, 2013). A previous study demonstrated that ROS can block the cell cycle progression *via* a FOXO-dependent inhibition on cyclin E/CDK2 (Owusu-Ansah *et al*., 2008). FOXO is a transcription factor that is also believed to play important roles in regulating stem cell quiescence and survival (Cheung and Rando, 2013). We found that FOXO protein level was markedly increased when either Complex III or Complex IV was disrupted (Figures 3C and 4A), which spurred us to test whether the ROS-FOXO pathway was involved in inhibiting ISC proliferation and differentiation in mutant midguts.

We first assessed ROS levels by staining midguts with CellROX (Sekihara *et al*., 2016; Manent *et al*., 2017), a fluorescent indicator for ROS. CellROX intensity drastically increased in guts pretreated with an organic peroxide, tert-butyl hydroperoxide, but markedly reduced in guts pretreated with a reducing agent, N-acetylcystine (Supplemental Figure S6, A and B), suggesting that CellROX indeed properly indicates the level of ROS. We found that intensities of CellROX were comparable between control and mutant midguts (Supplemental Figure S6, A and B). Additionally, co-overexpression of the ROS-scavenging enzymes SOD2 and catalase (Sun *et al*., 2002), did not rescue the differentiation phenotypes in ts mutant background (Supplemental Figure S6, C and D). These results indicate that ROS is not elevated in mutant cells, nor involved in the ISC differentiation phenotype. By contrast, FOXO protein was markedly increased in ISCs and EBs in mutant midgut compared to control (Figure 4A). To assess FOXO activity, we used a *lacZ* reporter under the control of the promoter of 4E-binding protein (4EBP) gene, a target of FOXO (Puig *et al*., 2003; Demontis and Perrimon, 2010). We found that lacZ protein was markedly higher in mutant cells than controls (Supplemental Figure S7). Taken together, these results demonstrate that FOXO protein and its transcriptional activity are elevated in mutant cells, in the absence of any detectable increase in cellular ROS.

**FIGURE 4:**
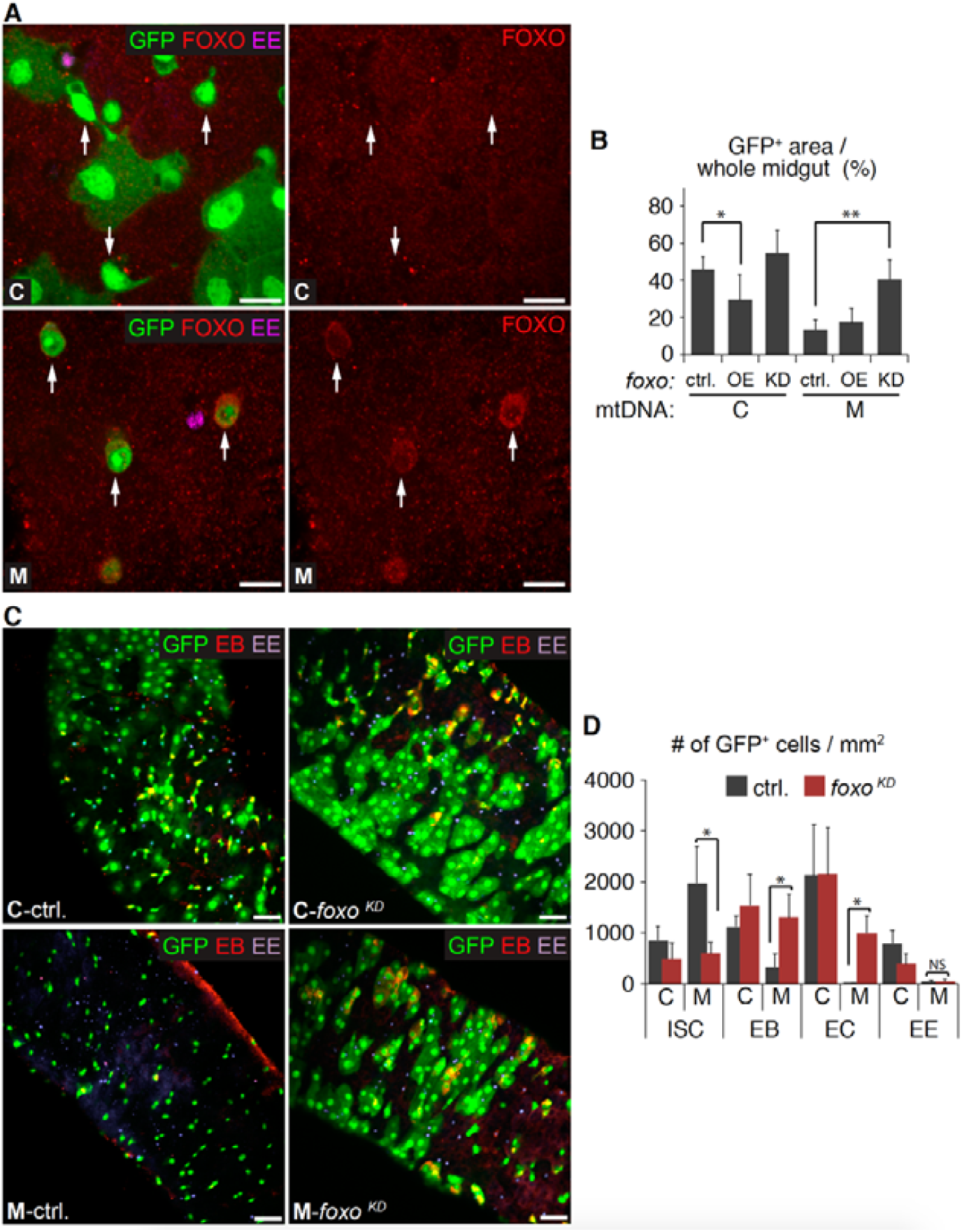
ETC disruption blocks ISC-to-EC differentiation through FOXO. (A) FOXO antibody staining of the control (C) and mutant (M) midguts (*esg*^*ts*^*F/O*>*mitoXhoI* flies), 10 days after the heat shock. FOXO protein levels dramatically increased in M compared to C ISCs/EBs (arrows). Green: GFP; Red: FOXO; Magenta: Prospero; Blue: DAPI. Scale bars: 10 µm. (B) Quantification shows that *foxo* overexpression (OE) in ISCs significantly reduced the GFP^+^ midgut area in the C group, 10 days after heat shock, while *foxo* knockdown (KD) significantly increased the GFP^+^ area in the M group. n=10 midguts each group, error bar: SD; *: p<0.01, **: p<0.001. (C) Views of the posterior parts of C and M midguts expressing a foxo knockdown construct (*esg*^*ts*^*F/O>mitoXhoI, foxo*^*KD*^ flies) or the RNAi control (*esg*^*ts*^*F/O>mitoXhoI, luciferase*^*KD*^ flies) 20 days after heat shock. (D) Quantification of C and M GFP^+^ midgut cell types in the presence or absence of foxo knockdown (*foxo*^*KD*^). *foxo*^*KD*^ led to dramatic increase in GFP^+^ EBs and ECs in M, but not GFP^+^ EEs. Green: GFP; Red: Su(H)GBE-lacZ; Purple: Prospero. Scale bars: 20 µm. n=10 midguts each group, error bar: SD; *: p<0.001, NS: Not Significant.

To understand the consequence of FOXO accumulation, we over-expressed FOXO using the *esg*^*ts*^*F/O* system in control midguts and found that it significantly reduced the GFP^+^ area (Supplemental Figure S8 and Figure 4B), indicating that increased FOXO activity can indeed block ISC differentiation. Reciprocally, knocking down *foxo* markedly increased the GFP^+^ area in mutant midguts (Supplemental Figure S8; Figure 4, B and C). Further classification of cell types demonstrated that *foxo* knockdown in ISCs or EBs significantly restored GFP^+^ EC but not GFP^+^ EE cell numbers (Figure 4, C and D). Altogether, our results suggest that FOXO protein is elevated in mutant midguts, which blocks the differentiation of ISCs to ECs.

### FOXO’s target Thor mediates the impairment in EC differentiation in mutant midguts

To determine how FOXO interferes with ISCs differentiation, we performed RNA sequencing (RNAseq) on control and mutant ISCs. About 10% of the mRNAs we detected were either up-regulated or down-regulated in the mutant ISCs relative to control (Supplemental Table S1). The down-regulated pool included many genes involved in cell differentiation, proliferation, and growth regulation (Supplemental Table S2). The up-regulated pool included genes participating in various metabolic processes (Supplemental Table S2). This result suggests a possible metabolic adaptation in response to the OXPHOS deficiency in mutant midguts. Importantly, knockdown of *foxo* in mutant ISCs restored the expression level of more than 50% of the up- or down-regulated genes (Figure 5A and Supplemental Table S2), indicating that transcriptional response to COX deficiency is partially mediated by FOXO.

**FIGURE 5:**
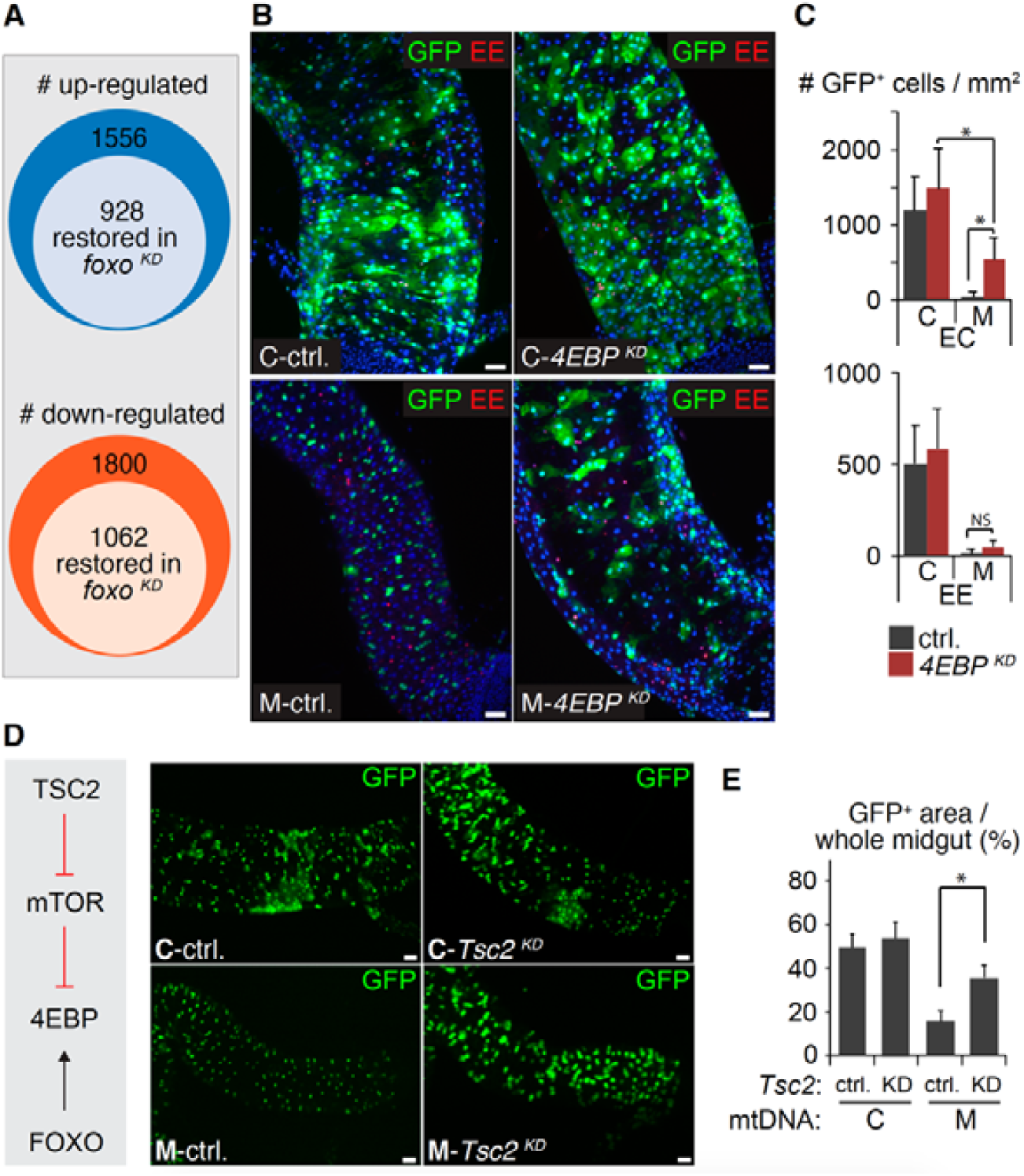
The impaired EC differentiation is partially mediated by 4EBP in mutant midguts. (A) Diagrams summarizing the number of genes up- or down-regulated by ETC disruption and restored by foxo knockdown (*foxo* ^*KD*^). The cut-off was 0.5 CPM fold change averaged across two replicas for regulation by ETC disruption, and 0.25 CPM fold change on average for restoration by *foxo* ^*KD*^. Full raw data is presented in Table S1. (B) Views of posterior midguts expressing a 4EBP knockdown construct (*esg*^*ts*^*F/O>mitoXhoI, 4EBP*^*KD*^ flies) compared to RNAi control (*esg*^*ts*^*F/O>mitoXhoI, luciferase*^*KD*^ flies) showed partial rescue of GFP^+^ EC numbers in M background. Green: GFP; Red: Prospero; Blue: DAPI. Scale bars: 20 µm. (C) Quantification of the GFP^+^ EC and EE cells in the posterior parts of C and M midguts in the presence or absence of *4EBP* ^*KD*^, 10 days after the heat shock. n=10 midguts each group, error bar: SD; *: p<0.001, NS: Not Significant. (D) Activation of mTOR pathway by Tsc2 knockdown in ISCs showed enlarged GFP^+^ cells in the M group midguts (*esg*^*ts*^*F/O>mitoXhoI, Tsc2* ^*KD*^ flies), indicating a partial rescue. Assessed 10 days after heat shock. Scale bars: 20 µm. (E) Quantification shows that *Tsc2* knockdown (KD) in ISCs significantly increased the GFP^+^ midgut area due to enlarged GFP^+^ cells in the M group, 10 days after heat shock. n=10 midguts each group, error bar: SD; *: p<0.001.

*Thor*, the sole homolog of *4EBP* in the *Drosophila* genome, was among the genes whose expression was upregulated in mutant midguts and restored by *foxo* knockdown (Supplemental Table S1). This finding is consistent with previous studies showing that *Thor* is a downstream target of FOXO in the regulation of cell growth and metabolism (Miron *et al*., 2001; Teleman *et al*., 2005). 4EBP proteins are highly conserved in metazoans, where they block the initiation of cap-dependent mRNA translation by binding to elongation factor eIF4E (Hernandez *et al*., 2010; Peter *et al*., 2015). The differentiation from EB to EC demands large amount of protein synthesis to drive cell growth, suggesting a mechanism by which the upregulation of Thor may block EB to EC differentiation in mutant midguts. To test this idea, we knocked down *Thor* in mutant EB cells using *esg*^*ts*^*F/O*-driven RNAi. We found that the number of GFP^+^ ECs was partially restored (Figure 5, B and C), confirming that the defect in EB-to-EC differentiation is partially mediated by Thor. In consistent with this observation, activating a 4EBP-inactivation pathway, the mTOR pathway, via *Tsc2* knockdown (Kapuria *et al*., 2012), leads to significantly enlarged GFP positive cells in mutant midguts (Figure 5, D and E), also indicating a partial rescue of the EC defect.

So far, our data indicate that ETC deficiencies increase FOXO protein level, which blocks ISC-to-EC differentiation through the 4EBP homolog Thor. However, foxo knockdown only restored the expression of a subset of genes altered in the mutant ISCs (Figure 5A and Supplemental Table S2). Additionally, neither *foxo* nor *thor* knockdown restored the number of GFP^+^ EE cells in mutant midguts (Figures 4D and 5C). These observations suggest that pathways other than foxo likely to be involved in regulating ISC-to-EE differentiation. Notch signaling is one of the major orchestrators of ISC differentiation and EE cell specification in the *Drosophila* midgut (Ohlstein and Spradling, 2007; Bardin *et al*., 2010; Biteau *et al*., 2011; Perdigoto and Bardin, 2013). We did notice that genes affected in mutant ISCs included many related to Notch signaling (Table S2). One of the AS-C complex gene, *scute (sc)*, which is negatively regulated by Notch signaling (Bardin *et al*., 2010), and required for EE specification (Amcheslavsky *et al*., 2014), was among the most severely down-regulated genes (Table S1). Additionally, the level of the intracellular domain of Notch, the active form of Notch appeared higher in mutant ISCs than control (Figure 6, A and B), indicating a mis-regulation of Notch in mutant ISCs. Noteworthy, mitochondrial disruption can enhance Notch signaling *via* elevated cytoplasmic Ca^2+^ level and calcineurin activity in mouse cardiomyocytes (Kasahara *et al*., 2013), suggesting a possible mechanism of Notch mis-regulation in mutant ISCs. Nonetheless, the exact mechanism and the pathophysiological consequences of this mis-regulation await future investigation.

**FIGURE 6:**
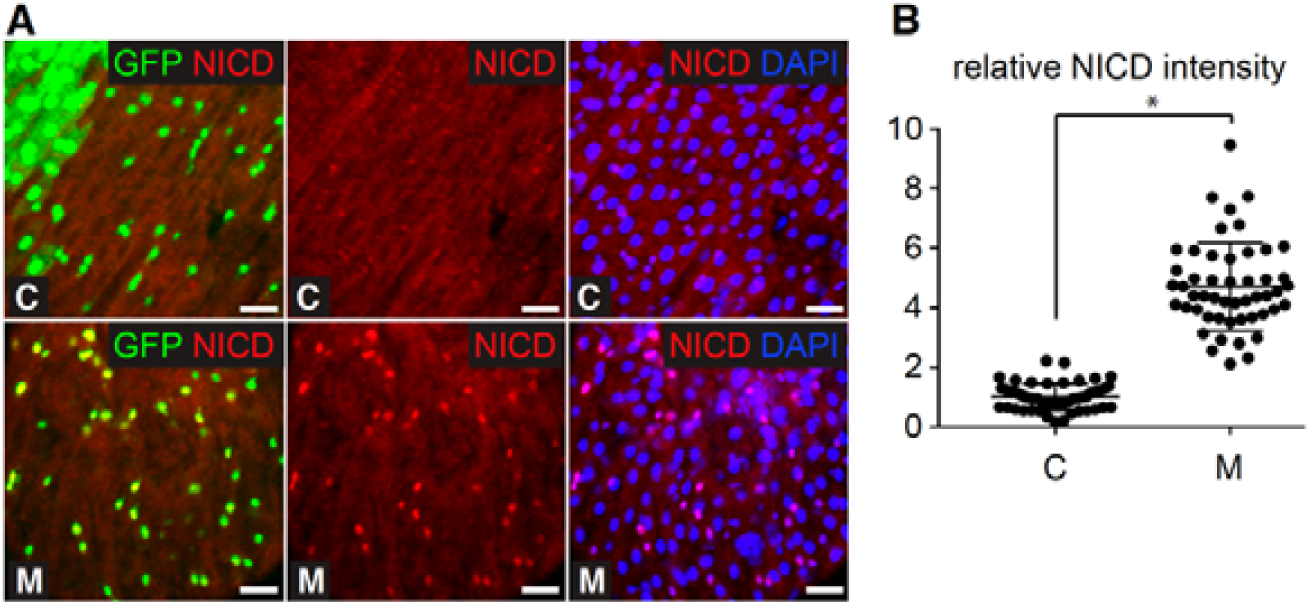
Nuclear NICD levels are elevated in mutant ISCs. (A) NICD antibody staining of midguts showing higher nuclear NICD levels in mutant (M) compared to control (C) ISCs (*esg*^*ts*^*F/O>mitoXhoI* flies). Green: GFP; Red: NICD; Blue: DAPI. Scale bars: 20 µm. (B) Quantification of the data as shown in (A). n=50, error bar: SD; *: p<0.001.

## DISCUSSION

While pharmacological disruptions of ETC complexes in cultured embryonic stem cells indicate that ETC activity is required for stem cell differentiation (Chung *et al*., 2007; Mandal *et al*., 2011), how it exerts this function is still enigmatic. Besides these experiments, carried out *in vitro* and further complicated by the drugs’ potential side effects, little evidence has been provided to clarify the role of ETC complexes in stem cell differentiation *in vivo*, especially for somatic stem cells maintaining tissue homeostasis during aging, such as ISCs. To address this question, we genetically disrupted ETC in ISCs by manipulating their mtDNA while keeping their environmental epithelial niche intact. We found that ETC deficiency in ISCs dramatically impaired their differentiation, though not necessarily because of ATP deficiency, the common consequence of ETC disruption, but by hyper-activating FOXO and mis-regulation of other signaling pathways.

Our conclusion that ATP deficiency is not the main driver of differentiation impairment in our mutant ISCs stems from two observations: 1) knocking down individual OXPHOS complexes with RNAi did not always phenocopy the defects observed with our mutant (Figure 3B; Supplemental Figure S5, B-D); 2) knocking down AMPK, which is activated by ATP deficiency, failed to rescue the phenotypes of the *ts* mutant (Supplemental Figure S4, B and C). In our system, disruption of Complexes I, II and V (mitochondrial ATP synthase) with RNAi against nuclear-encoded subunits had little effect on ISC differentiation. By contrast, disruption of Complexes III and IV nuclear-encoded subunits resulted in the similar differentiation defects as our mutant. While perplexing, these observations are not unique, as a previous study (Teixeira, Sanchez, Hurd, *et al*., 2015), using the same RNAi constructs, showed that the differentiation of Drosophila female germ line stem cells was independent of ATP production. Moreover, *AMPK* knockdown has been shown to rescue cell proliferation defect caused by an nuclear COX mutation in the *Drosophila* eye (Mandal, Guptan, *et al*., 2005), however, its inability to rescue the ISC differentiation in our mutant supports the conclusion that, at least in ISCs, ETC promotes differentiation through ATP-independent pathways.

Another perplexing observation is that, in our system, disruption of ETC does not lead to ROS accumulation (Supplemental Figure S6, A and B). This stands in contrast to reports that leakage of free electrons to oxygen between Complexes III and IV is a main source of cellular ROS (Starkov, 2008; Bleier and Drose, 2013). Importantly, overexpression of ROS scavengers did not rescue the mutant phenotype (Supplemental Figure S6, C and D), further indicating that ROS accumulation is not the cause of the differentiation defect. Yet, we find that our mutant leads to elevated expression of FOXO (Figure 4A), a transcription factor known to be upregulated in response to ROS stress and mediate stem cell behavior (Cheung and Rando, 2013; Klotz *et al*., 2015). How ETC disruption leads to elevated FOXO activity in our system (Figure 4A and Supplemental Figure S7) is currently unclear. Hyper-activation of FOXO and elevated FOXO levels have been documented in human disease accompanied with mitochondrial dysfunction and increased oxidative stress (Kannike *et al*., 2014). The increased FOXO protein levels in these contexts result from FOXO autoregulation via a positive feedback loop. Such positive auto-feedback loop is also reported in the Drosophila adult gut (Alic *et al*., 2014), and may remain active in the mutant ISCs. Aside from ROS, the mitochondrial unfolded protein response (UPR^mt^) causes lasting FOXO activation during *Drosophila* development (Borch Jensen *et al*., 2017). Could the reduced availability of mtDNA-encoded subunits in mutant ISCs result in UPR^mt^ and eventually activate FOXO? In this case, the impact of an ETC deficiency on ISC differentiation could be indirect. However, it is also possible that some ETC complexes play specific roles besides respiration. For instance, a recent study of Parkinson’s disease in *Drosophila* model (Meng, Yamashita, *et al*., 2017), led to the proposal that Complex III and IV help maintain mitochondrial crista structure. The possibility of a specific impact of Complexes III and IV on ISC differentiation can therefore not be eliminated, though the exact mechanisms remain obscure at this point.

As for the impact of elevated FOXO on the ISC lineage, several scenarios are possible. *Drosophila* FOXO functions as a key negative regulator of growth and proliferation, probably via 4EBP transcription (Puig *et al*., 2003). Indeed, we found that over-expression of FOXO in normal ISCs blocked their proliferation and differentiation (Supplemental Figure S8 and Figure 4B), and inhibition of FOXO in mutant ISCs partially rescued the defect (Figure 4, C and D), as did down-regulation of 4EBP (Figure 5, B and C). Considering the large size of EC cells that demand prodigious protein synthesis, it is not far-fetched to propose that activation of the FOXO-4EBP cascade should inhibit the formation of EC cells through its inhibition on protein translation. Additional support for this notion stems from our observation that activating mTOR, a protein kinase that inactivates 4EBP, by knocking down its upstream inhibitor Tsc2 (Kapuria *et al*., 2012), leads to significantly enlarged GFP positive cells in mutant midguts, indicating a partial rescue of the EC defect (Figure 5, D and E). Thus, we conclude that FOXO acts at least in part by blocking the growth of ECs through 4EBP. Besides cell growth, FOXO also regulates metabolism (Nirala, Rahman, *et al*., 2013), and its upregulation in the ts mutant may cause a shift in ISCs’ metabolism. Consistent with this hypothesis, Lactate dehydrogenase that converts pyruvate to lactate and thus shifts the energy metabolism away from OXPHOS (Wang *et al*., 2016), was significantly upregulated in mutant ISCs, but restored to normal levels by *foxo* knockdown (Supplemental Table S1). These results suggest that OXPHOS disruption in ISCs caused a metabolic shift that was partially contributed by elevated FOXO, and further support our finding that the intact ETC and aerobic metabolism are essential for ISCs’ activity and midgut epithelial homeostasis.

## MATERIALS AND METHODS

### Fly stocks and transgenic lines

Fly strains used in this study including the follows: *w;esg-GAL4,tub-GAL80*^*TS*^,*UAS-nlsGFP/CyO;UAS-flp,Act>CD2>GAL4/TM6B*(Jiang *et al*., 2009); *Su(H)GBE-LacZ* (Micchelli and Perrimon, 2006); *If/CyO;UAS-mitoXhoI; UAS-mitoXhoI /CyO;MRKS/TM6B; UAS-SOD2, Catalase; AMPKα* RNAi (BDSC#25931); *foxo* RNAi (BDSC #27656); *UAS-FOXO* (BDSC #42221); *ND30* (30kD) RNAi (VDRC #103412); *NDB12* (B12) RNAi (VDRC #104890); *ND23 (TYKY)* RNAi (VDRC #21748); *SdhA* RNAi (VDRC #110440); *SdhC* RNAi (VDRC #6031); *SdhD* RNAi (VDRC #101739); *Cytochorme c1(cyt c1)* RNAi (VDRC #109809); *UQCR-C1(cp1)* RNAi (VDRC #101350); *UQCR-C2(cp2)* RNAi (VDRC #100818); *COX5A(Va)* RNAi (VDRC #109070); *COX7C(VIIc)* RNAi (VDRC #104970); *COX8(VIII)* RNAi (VDRC #104047); *ATPsynB (b)* RNAi (VDRC #106758); *ATPsynC (c)* RNAi (VDRC #106834); *ATPsynD (d)* RNAi (VDRC #104353); *Thor-lacZ* (4EBP-lacZ, BDSC #9558); *Thor* RNAi (BDSC #36815); *Tsc2* RNAi (BDSC #35401); *UAS-pAbp* (BDSC #9420, For tissue specific mRNA tagging(Yang *et al*., 2005)); *Luciferase* RNAi (BDSC #31603, control for RNAi lines). Female flies of heteroplasmic *mt:CoI*^*T300I*^ and homoplasmic *mt:CoI*^*syn*^ background as previously described (Xu *et al*., 2008; Chen *et al*., 2015) were used to generate the mutant (M) group and control (C) group flies for assessments, respectively. All flies and crosses were maintained at 18 °C.

### Tissue preparation, immunofluorescence staining and confocal imaging

All fly crosses and the offspring were kept at 18 °C. Two days after eclosion, the adult flies for assay were heat-shocked at 37 °C for two hours and maintained at 29 °C until dissection at different day points. The entire midgut tissues were dissected out in room temperature Schneider’s medium (Thermo Fisher Scientific) supplied with 10% heat inactivated Fetal Bovine Serum (FBS, Thermo Fisher Scientific). The midguts were then rinsed by same medium and either used for directly imaging or further fixation and staining.

Immunofluorescence staining was done after tissue fixation in PBS containing 4% PFA and three times PBS washing. The antibodies and reagents used for staining were: mouse anti-Prospero (MR1A, Developmental Studies Hybridoma Bank, DSHB); mouse anti-Delta (C594.9B, DSHB); chicken anti-Beta Galactosidase (ab9361, Abcam); rabbit anti-Pdm1 (gift from Cai Yu); rabbit anti-Cleaved Caspase-3 (9661, Cell signaling); rabbit anti-FOXO antibody (ab195977, Abcam); GFP Tag Polyclonal Antibody, Alexa Fluor 488; Alexa Fluor secondary antibodies (Thermo Fisher Scientific) and Vectashield mounting medium with DAPI (Vector Laboratories).

Imaging analysis was performed using the Carl Zeiss Axio Observer equipped with Perkin Elmer spinning disk confocal system. For live imaging, a Zeiss incubation systems were used to maintain proper temperature, humidity and CO_2_ level. The image processing and quantification were performed by Volocity 6.1.1 software (Perkin Elmer) and Image J software (NIH), 10 midguts were analyzed per group. The region of the posterior midgut ∼500 μm from the junction of the midgut and hindgut was used for the quantification of GFP^+^ cell numbers.

### Detection of mitochondrial membrane potential

Mitochondrial membrane potential was detected using protocol adopted from a previously study (Zhang *et al*., 2019). Briefly, after dissection, intact guts were incubated in the Schneider’s medium containing TMRM (200nM, Thermo Fisher Scientific) and MitoTracker Deep Red (200nM, Thermo Fisher Scientific) at 29°C for 20 min, rinsed with medium for 3 times. Samples from different groups were mounted with medium on the same coverslip set in a custom-made metal frame and then covered with a small piece of Saranwrap before imaging. Images were captured with fixed confocal settings. These small, isolated and basally-localized GFP^+^ cells within the intestinal epithelium were identified as ISCs in esg^ts^ GFP flies based on a previously study (Hochmuth *et al*., 2011). A single section of basal epithelium was captured, and TMRM and MitoTracker signal intensities were quantified using Image J. Mitochondrial membrane potential was computed as the ratio of the mean intensity of TMRM to MitoTracker fluorescence with background correction.

### ATP measurement

Tissue ATP level was measured using the ATP Bioluminescence Assay Kit HS II (Roche) following the manufacturer’s protocol. Wild-type and homoplasmic ts female flies were raised at 18°C for 2 days after eclosion and then shifted to 29°C for 3 days, or continuously maintained at 18°C as control. For each genotype, individual midgut was dissected out and immediately homogenized in 100μL lysis buffer. The lysates were boiled for 5 min and centrifuged at 20,000 g for 1 min. 10μL cleared lysate was used for ATP measurement using the Kit through a 96-well plate luminometer (Berthold). The amount of ATP was then determined and normalized with the protein level measured by BCA protein assay (Thermo Fisher Scientific).

### Detection of ROS levels

ROS levels were detected using CellROX Deep Red following the manufacturer’s protocol (Thermo Fisher Scientific). Briefly, the guts were incubated in the Schneider’s medium containing 5 μM CellROX Deep Red for 45 minutes at 29°C followed by three washes with PBS. The samples were then fixed with 3.7% formaldehyde for 15 minutes and mounted for imaging analysis. For comparison and quantification, the dissected guts were pre-incubated in the Schneider’s medium with or without drugs for one hour at 29°C before CellROX staining. Control group guts incubated in the medium containing 200 μM tert-butyl hydroperoxide (TBHP) worked as the positive controls, and incubated with 1mM N-acetylcysteine (NAC) as the negative controls. Images were captured with a spinning disc confocal. These small, isolated and basally-localized GFP^+^ cells within the intestinal epithelium were identified as ISCs in esg^ts^ GFP flies based on a previously study (Hochmuth et al., 2011). A single section of basal epithelium was captured, and CellROX intensity was quantified using Volocity 6.1.1 software (Perkin Elmer).

### GFP positive ISCs sorting and mtDNA level

The GFP^+^ ISCs is sorted as previously described (Dutta *et al*., 2013). Briefly, midguts of “w; *esg-GAL4, tub-GAL80*^*ts*^ *UAS-nlsGFP/UAS-mitoXhoI”* flies with different mtDNA background, which was heat-shocked and maintained in 29 °C for 7 or 15 days, were dissected in PBS and dissociated for 2 h by incubating with 0.8 mg/ml elastase at 29°C. ISCs were then sorted based on intensity of GFP marker under the help of Flow Cytometry (FACS) Core Facility of NHLBI, NIH. To quantify mtDNA level, total DNA was prepared from the sorted samples using the DNeasy Blood kit (Qiagen). Quantitative Real-time PCRs were performed in triplicate using Roche LightCycler 480 system and SYBR Green I Master. Primers based on mtCol (F: ATTGGAGTTAATTTAACATTTTTTCCTCA; R: AGTTGATACAATATTTCATGTTGT-GTAAG) and His4 (F: TCCAAGGTATCACGAAGCC; R: AACCTTCAGAACGCCAC) sequences was used for the mtDNA and nuDNA, respectively.

### EdU Incorporation

EdU Incorporation was done following the as previously described (Zhang *et al*., 2015), using the Click-iT EdU Alexa Fluor 555 Imaging kit (Thermo Fisher Scientific). Briefly, intact guts were dissected from adult female flies, and were incubated in 10 μM EdU in the Schneider’s medium containing 10% FBS for 2 h at 25°C. Tissues were then fixed with 4% PFA, washed in PBS with 0.1% Triton X-100 (PBST), processed with the Kit and mounted for imaging analysis.

### ISC specific mRNA profiling

A modified mRNA tagging technique (Yang *et al*., 2005) is used to isolate mRNA from ISCs of fly midguts. Briefly, the FLAG-tagged poly(A)-binding protein (pAbp) is expressed in C and M group flies under esg-GAL4 driver. About 200 midguts from C and M group “*esg-GAL4,tub-GAL80*^*TS*^,*UAS-nlsGFP/UAS-mitoXhoI;UAS-pAbp-FLAG/+*” or “*esg-GAL4,tub-GAL80*^*TS*^,*UAS-nlsGFP/UAS-mitoXhoI;UAS-pAbp-FLAG/UAS-FOXO-IR*” flies, which were heat-shocked and maintained in 29 °C for 5 days, were fixed in 1 ml of PBS containing 1% formaldehyde (Sigma-Aldrich) and 0.5% Igepal CA-630 (Sigma-Aldrich) for 1 h on ice, then, incubated for 5 min after addition of 140 μl of 2 M glycine. Samples were rinsed three times with ice cold PBS and homogenized by sonication in 2 ml of homogenization buffer (150 mM NaCl, 50 mM HEPES buffer at pH 7.6, 1 mM EGTA, 15 mM EDTA and 10% glycerol) supplied with 8 mM vanadyl ribonucleoside complex (Sigma-Aldrich), 50 U/ml SUPERase-In (Thermo Fisher Scientific), and 1 tablet/50 ml protease inhibitor cocktail tablet (Roche). After centrifugation at 13,000 g for 30 min, the supernatant was incubated with anti-FLAG-M2 affinity agarose beads (Thermo Fisher Scientific) at 4°C O/N. The beads were then washed four times with homogenization buffer. And the RNAs bound to the pAbp were recovered in 100 ml of elution buffer (50 mM Tris–HCl, pH 7.0, 10 mM EDTA, 1.3% SDS and 50 U/ml SUPERase-In) by proteinase K (40 μg/mL) treatment 2h at 60°C and O/N at 65°C after addition of NaCl (200mM). Eluant was then mixed with 4V of Trizol (Thermo Fisher Scientific) and 1V of chloroform (Sigma-Aldrich) with vigorously shaking, incubated at room temperature for 10 min, and then centrifuged at 12,000 g for 15 min at 4°C. The supernatant was transferred and mix with 2.5V of isopropanol, chilled at −80°C O/N, and centrifuged at 12,000 g for 30 min at 4°C. The RNA pellet was rinsed twice with 70% ethanol and then dissolved in RNase free water. Poly(A) capture libraries were generated at the DNA Sequencing and Genomics Core, NHLBI, NIH. RNA sequencing was performed with using a Hiseq3000 (Illumina) and 75-bp pair-end reads were generated at the DNA Sequencing and Genomics Core, NHLBI, NIH. RNA sequencing data were analyzed by the Bioinformatics and Computational Biology Core, NHLBI, NIH. After quality control (QC) assessment of FASTQ files using FastQC toolkit (v0.11.5) with default parameters. paired-end reads were aligned against Drosophila Melanogaster reference genome (Dmel6) using HISAT2 (v2.0.5) alignment. Gene level read counts were produced by featureCounts (v1.5.2) using paired-end, reversely stranded read. Differential expression analysis at the gene-level was carried out using limma-voom open source R packages. TMM normalization was carried out and normalized factors were estimated for each sample. The lmFit function in limma-voom was used to Fit linear models for each gene to calculate log2 fold changes and p-values using the normalized factors as weights in the model. The cut-off for gene up- or down-regulation in M relative to C was 0.5 CPM fold change averaged across two replicas, and the cut-off for restoration by *foxo* ^*KD*^ was 0.25 CPM fold change in M-*foxo* ^*KD*^ relative to C-*foxo* ^*KD*^.

The RNAseq data were deposited in Gene Expression Omnibus of NCBI and will be available with accession number (GEO: GSE137231, Secure token: onolokmgbrmzlgp) https://www.ncbi.nlm.nih.gov/geo/query/acc.cgi?acc=GSE137231

## Statistical analysis

F-test was performed to evaluate the equality of variances. Two-tailed Student’s t-test was used for statistical analysis. Difference was considered statistically significant when *P*<0.05. Results are represented as mean ± SD of the number of determinations.

## ACKNOWLEDGEMENT

We thank F. Chanut for comments and editing on the manuscript; B. Edgar, N. Perrimon, the Bloomington Drosophila Stock Center and the Vienna Drosophila Resource Center for various fly lines; Y. Cai and the Developmental Studies of Hybridoma Bank for various antibodies; the NHLBI DNA Sequencing and Genomics Core, and NHLBI FACS core for technical assistance. This work is supported by NHLBI Intramural Research Program.

## Supplemental Figures and Figure Legends

**Figure S1:**
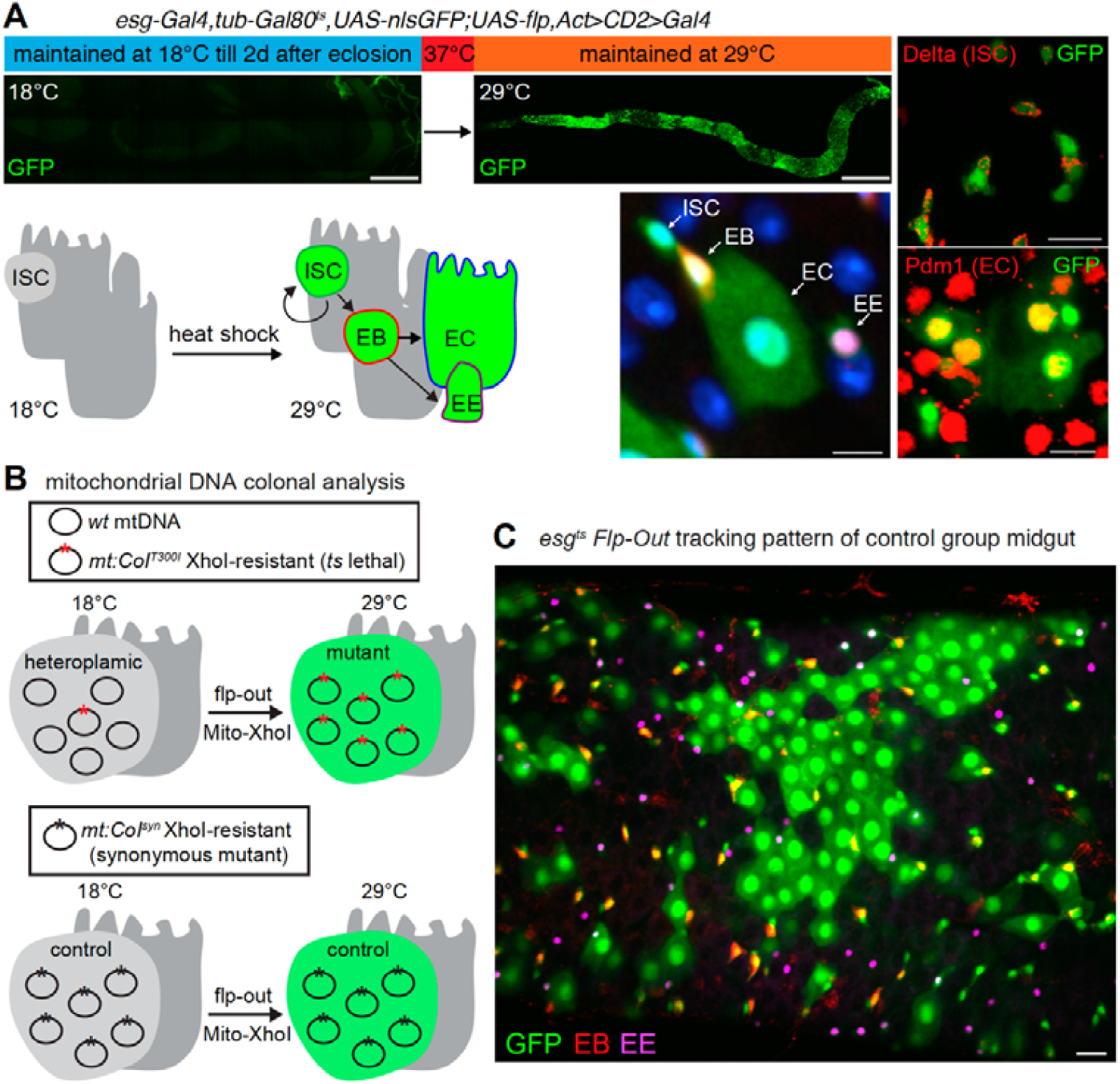
Strategy for studying the role of mitochondrial ETC system in intestinal stem cells. (A) Tracking ISCs with the *esg*^*ts*^ *Flp-Out* (*esg*^*ts*^*F/O*) labelling system. (Upper) “*esg*^*ts*^*F/O* “flies (i.e., *w*; *esg-GAL4, tub-GAL80*^*ts*^ *UAS-nlsGFP; UAS-flp, act>CD2>GAL4*) were grown at 18 ºC, exposed to a 2-hr heat shock at 37 ºC two days after eclosion into adults, and maintained at 29 ºC until assessment. Representative images show no GFP expression in the midgut of *esg*^*ts*^*F/O* adults before heat shock and a large percentage of GFP^+^ cells 20 days after heat shock. (Lower) Schematics of ISC labelling: heat shock and maintenance at 29 ºC reverse the inhibition of the GAL4/UAS expression system by GAL80^ts^, leading to continuous expression of GFP in esg^+^ ISCs and their lineage via Flp activation of Act-GAL4, without affecting other already differentiated midgut epithelial cells. Differentiation of the GFP^+^ ISCs into EBs, ECs and EEs can be tracked based on cell shape and the expression of specific cell markers. Green: GFP; Red: Delta antibody staining for ISC, Su(H)GBE-lacZ for EB and Pdm1 antibody staining for EC; Magenta: Prospero antibody staining for EE; Blue: DAPI. Scale bars: 200 µm for whole-gut views, 10 µm for close-up views. (B) Mitochondrial DNA clonal analysis. Wild type (wt) mtDNA carries a unique restriction XhoI site in the cytochrome c oxidase locus, *mt:CoI*. The *mt:CoI*^*T300I*^ mutation is resistant to XhoI cleavage but disrupts COXI function at high temperature (temperature-sensitive lethal). The *mt:CoI*^*syn*^ mutant mtDNA is also resistant to XhoI cleavage due to a synonymous mutation that does not affect COXI function. Using the *esg*^*ts*^*F/O* system, we can drive the expression of a mitochondria-targeted form of XhoI (*mitoXhoI*) in the ISCs of flies carrying either a heteroplasmic mixture of wt and *mt:CoI*^*T300I*^ mtDNAs (mutant group), or a homoplasmic population of *mt:CoI*^*syn*^ mtDNA (control group). The heteroplasmic mixture is more than 80% WT and behaves as wild type even at 29 ºC. After heat shock and maintenance at 29 ºC, WT mtDNAs are eliminated by mitoXhoI cleavage in ISCs only—the ECs and EEs that formed before the heat shock do not express *esg* and remain heteroplasmic. The control group carrying the homoplasmic *mt:CoI*^*syn*^ mutation, which is resistant to mitoXhoI cleavage, retains wild-type COXI function at both 18 ºC and 29 ºC. This method allows the disruption of ETC system in the GFP^+^ ISC cells without affecting the GFP^-^ midgut epithelial tissues. (C) Representative *esg*^*ts*^ *F/O* tracking pattern in the midgut of control flies, showing patches of GFP^+^ EBs, EEs and ECs over a background of non GFP tissues, reflecting past ISC proliferation/differentiation events. Green: GFP; Red: Su(H)GBE-lacZ; Magenta: Prospero. Scale bar: 10 µm.

**Figure S2:**
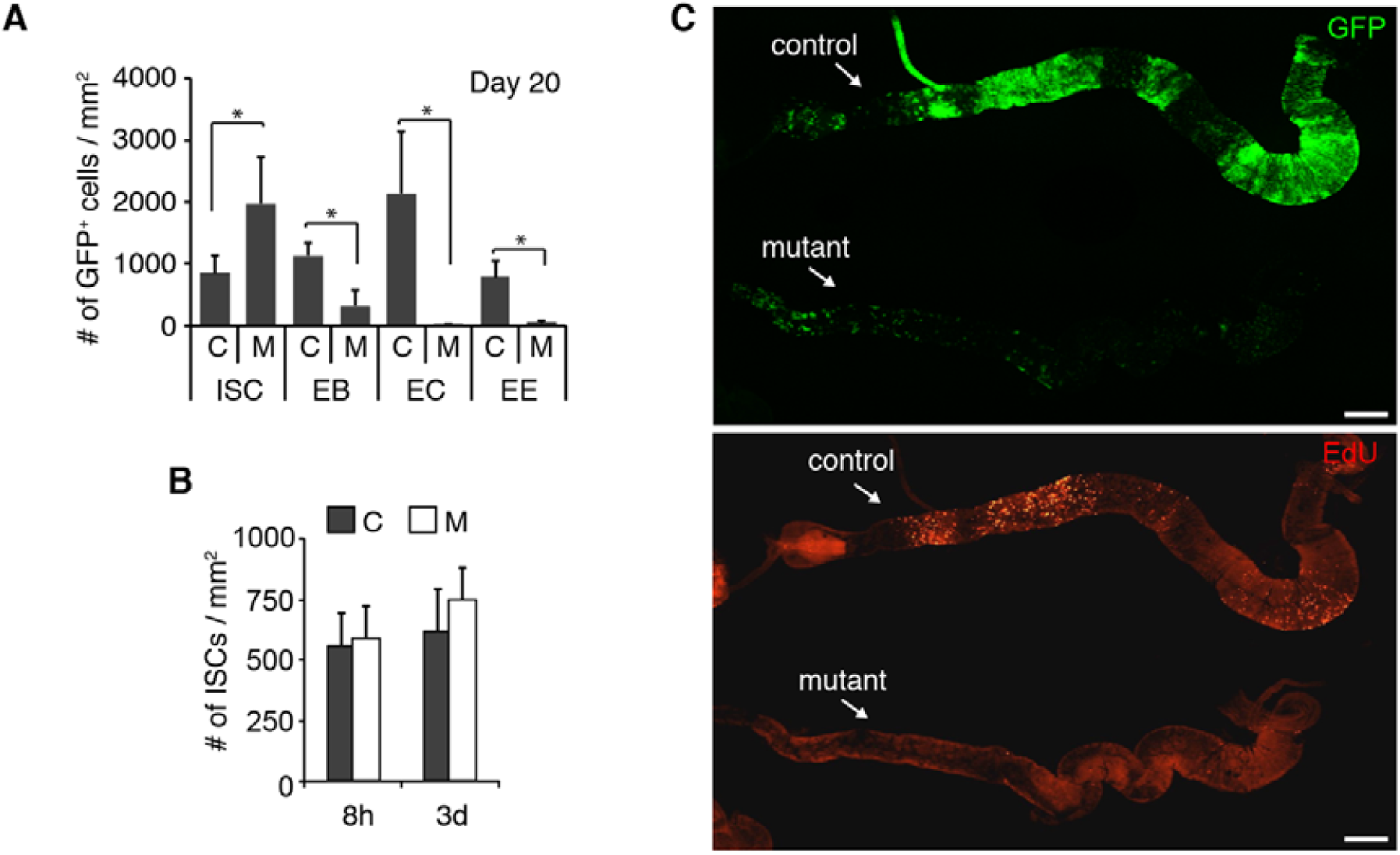
Less GFP^+^ differentiated cells and amount of EdU incorporation in M group. (A) Quantification of the GFP^+^ ISCs, EBs, ECs and EEs in the posterior parts of the control (C) and mutant (M) midguts, 20 days after the heat shock. n=10 midguts each group, error bar: SD; *: p<0.001. (B) Quantification of ISCs in C and M anterior midguts of the “*esg*^*ts*^*>mitoXhoI*” flies, 8 hour and 3 days after heat shock. n=10 midguts each group, error bar: SD. (C) EdU incorporation assay of C and M midguts, 20 days after the heat shock. The representative C and M midguts were assayed together and imaged in the same view. EdU signals are dramatically reduced in the M midgut. Green: GFP; Red: EdU. Scale bars: 200 µm.

**Figure S3:**
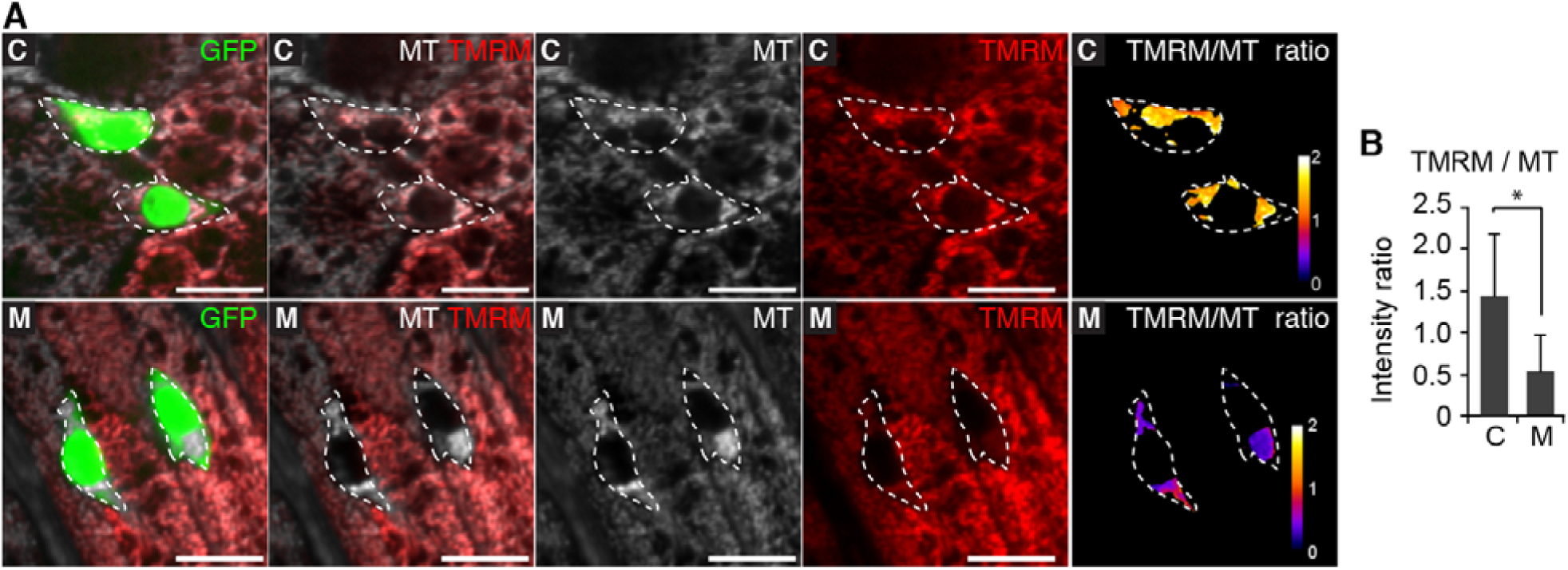
Impaired mitochondrial membrane potential in mutant ISCs. (A) Representative views of midguts stained with TMRM (a membrane potential marker) and MitoTracker Deep Red (MT, a mitochondrial mass marker), of the control (C) and mutant (M) group “*esg*^*ts*^*>mitoXhoI*” flies, 10 days after the heat shock. ISCs are indicated by dash lines. Green: GFP; Gray: MitoTracker Deep Red; Red: TMRM; ratiometric image: TMRM/MT. Scale bars: 10 µm. (B) Quantification of the ratios of TMRM to MitoTracker Deep Red (MT) in ISCs of C and M “*esg*^*ts*^*>mitoXhoI*” flies, n=50 total ISCs, 10 midguts each group, error bar: SD; *: p<0.001.

**Figure S4:**
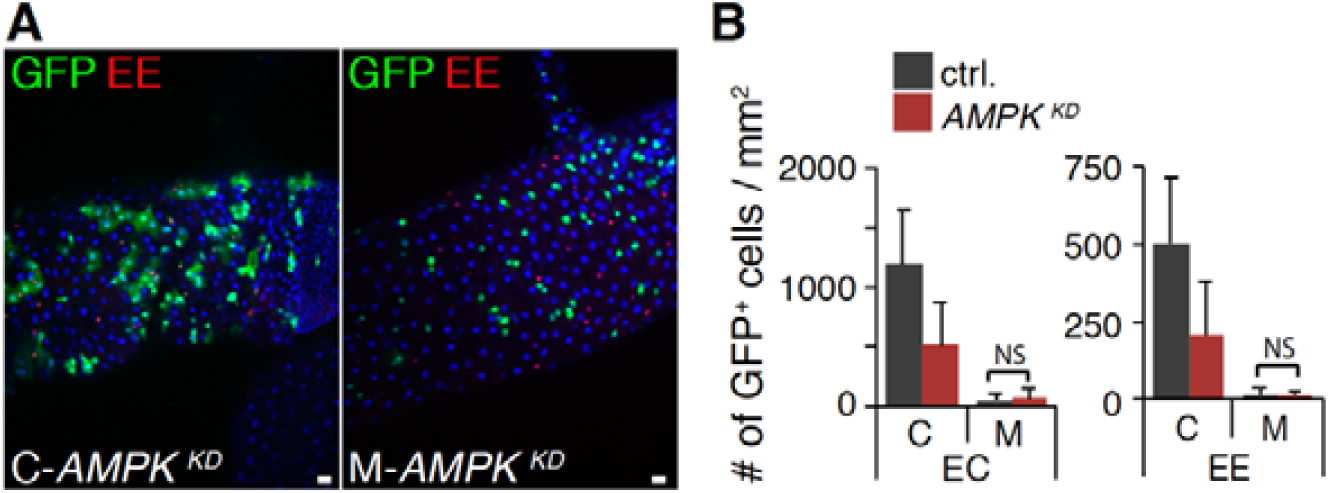
AMPK knockdown did not rescue the M group midgut phenotype. (A) The *AMPK* knockdown (*AMPK*^*KD*^) did not rescue the M group midgut phenotype in “*esg*^*ts*^*F/O>mitoXhoI*” files, 10 days after heat shock. Scale bars: 10 µm. (B) Quantification of the GFP^+^EC and EE cells in the posterior parts of C and M midguts expressing a AMPK knockdown construct (*esg*^*ts*^*F/O>mitoXhoI, AMPK* ^*KD*^ flies), or a RNAi control (*esg*^*ts*^*F/O>mitoXhoI, luciferase*^*KD*^ flies), 10 days after the heat shock. n=10 midguts each group, error bar: SD; NS: Not Significant.

**Figure S5:**
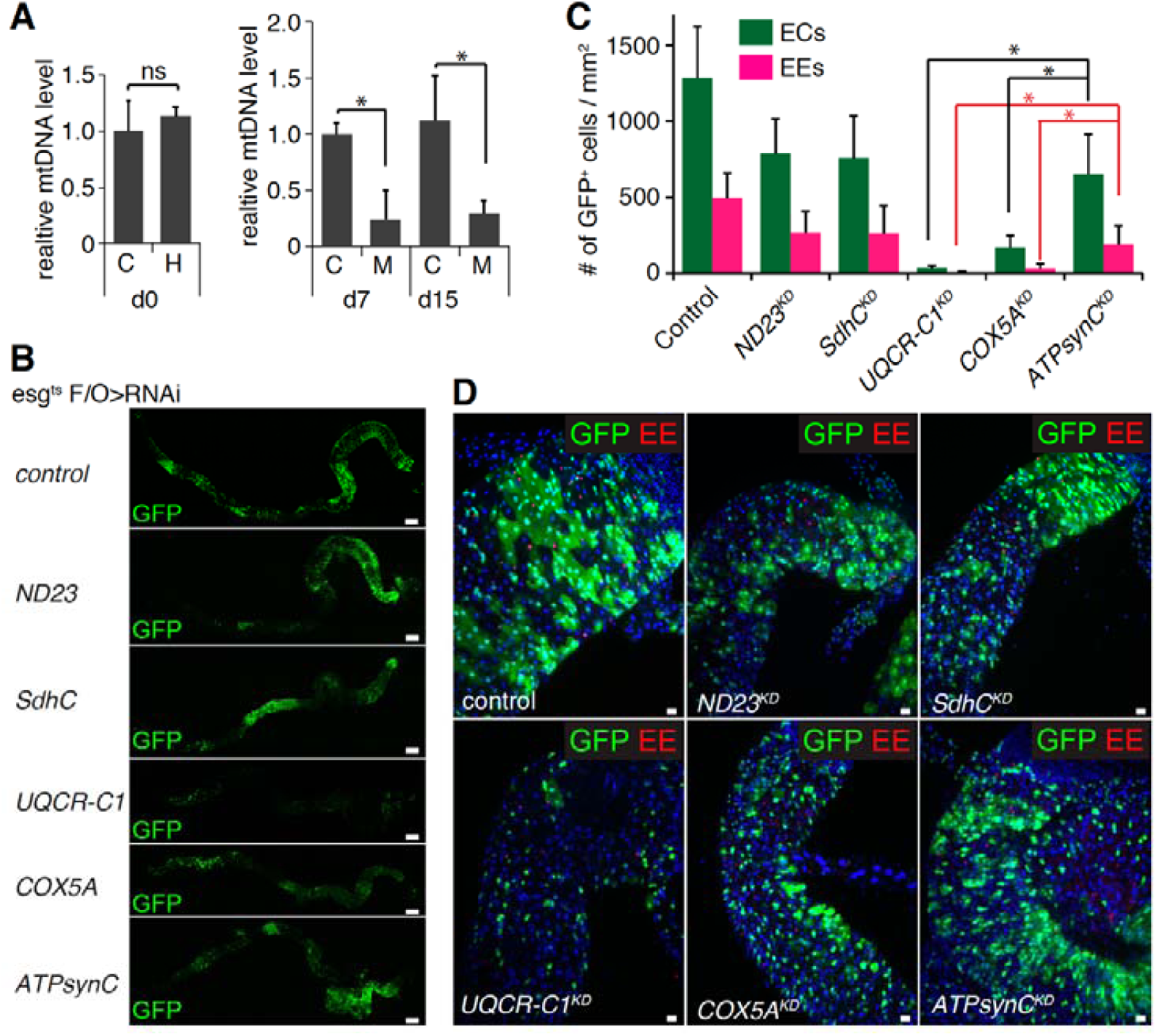
ETC disruption blocks ISC differentiation via pathways independent of mitochondrial energetic deficiency. (A) Real-time PCR shows that, before heat shock, there is no significant difference in the mtDNA levels between the control (C) and heteroplasmic (H) midguts of “*esg*^*ts*^*>mitoXhoI*” flies. However, after the heat shock, there are significantly reduced mtDNA levels in the sorted GFP^+^ISCs of mutant (M) group “*esg*^*ts*^*>mitoXhoI*” flies when compared with C group at day 7 and 15. n=3, error bar: SD; *: p<0.001, ns: Not Significant. (B) Knockdown of the key subunit of each respiratory complex in the ISCs of “*esg*^*ts*^*F/O*” flies reveals that disruption of the UQCR-C1 (TYKY, Complex III) or COX5A (Va, Complex IV) function leads to the most severe ISC differentiation defect, 10 days after the heat shock. Green: GFP. Scale bars: 200 µm. (C) Quantification of the GFP^+^EC and EE cells in the midguts of “*esg*^*ts*^*F/O*” flies combined with individual knockdown (KD) of the respiratory complex subunit, 10 days after heat shock. (D) Both GFP^+^EC and EE cells are dramatically reduced in the midguts of “*esg*^*ts*^*F/O>UQCR-C1* ^*KD*^” or “*esg*^*ts*^*F/O>COX5A* ^*KD*^” flies, 10 days after heat shock. The ISC differentiation events are almost missing in the midguts of “*esg*^*ts*^*F/O>UQCR-C1*^*KD*^”. Green: GFP; Red: Prospero; Blue: DAPI. Scale bars: 10 µm.

**Figure S6:**
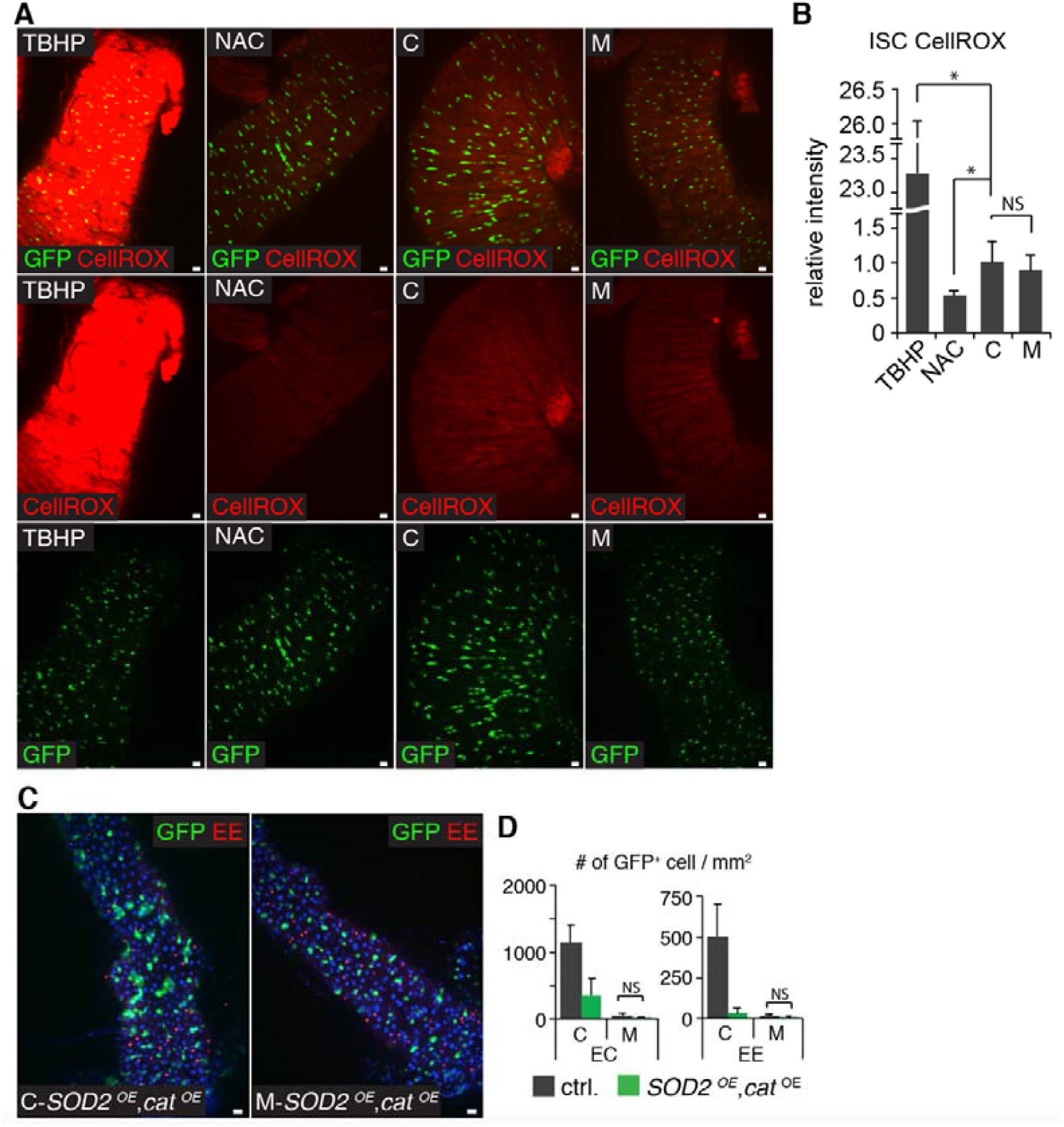
ROS is not elevated in the mutant ISCs and ROS scavengers do not rescue the mutant phenotype. (A) CellROX Deep Red (CellROX) staining of the control (C) and mutant (M) group midguts of the “*esg*^*ts*^*>mitoXhoI*” flies, 10 days after the heat shock. Tert-butyl hydroperoxide (TBHP) treatment of C group midguts were used as the positive controls, and N-acetylcysteine (NAC) treatment of C group midguts were used as the negative controls. Scale bars: 10 µm. (B) Quantification of the CellROX intensities as in (A) relative to C group ISCs, n=50 total ISCs, 10 midguts each group, error bar: SD; *: p<0.001, NS: Not Significant. (C) The co-overexpression (OE) of two ROS scavengers, SOD2 and Catalase did not rescue the mutant phenotype in the M group midgut of the “*esg*^*ts*^*F/O>mitoXhoI*” flies, 10 days after heat shock. There was no rescue of GFP^+^ EC and EE cell numbers in M group but rather reduced GFP^+^ EC and EE numbers in C group. Green: GFP; Red: Prospero; Blue: DAPI. Scale bars: 10 µm. (D) Quantification of the GFP^+^EC and EE cells in the posterior parts of C and M midguts overexpressing SOD2 plus Catalase (*esg*^*ts*^*F/O>mitoXhoI, SOD2*^*OE*^,*Catalase* ^*OE*^ flies), 10 days after the heat shock. n=10 midguts each group, error bar: SD; NS: Not Significant.

**Figure S7.**
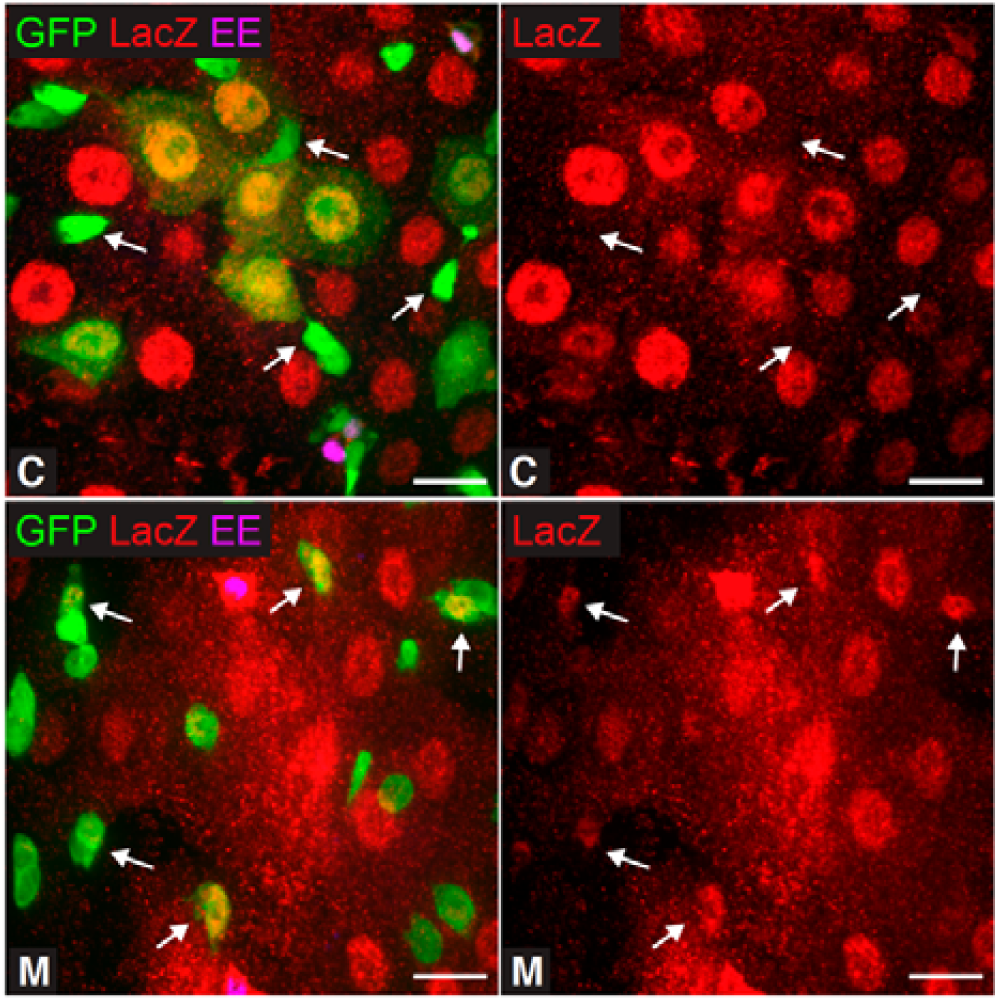
FOXO activity is elevated in mutant ISCs/EBs. FOXO activity in C and M midguts as revealed by 4EBP::lacZ expression. FOXO activity dramatically increased in M compared to C ISCs/EBs (arrows), 10 days after the heat shock. Green: GFP; Red: 4EBP::lacZ; Magenta: Prospero. Scale bars: 10 µm.

**Figure S8.**
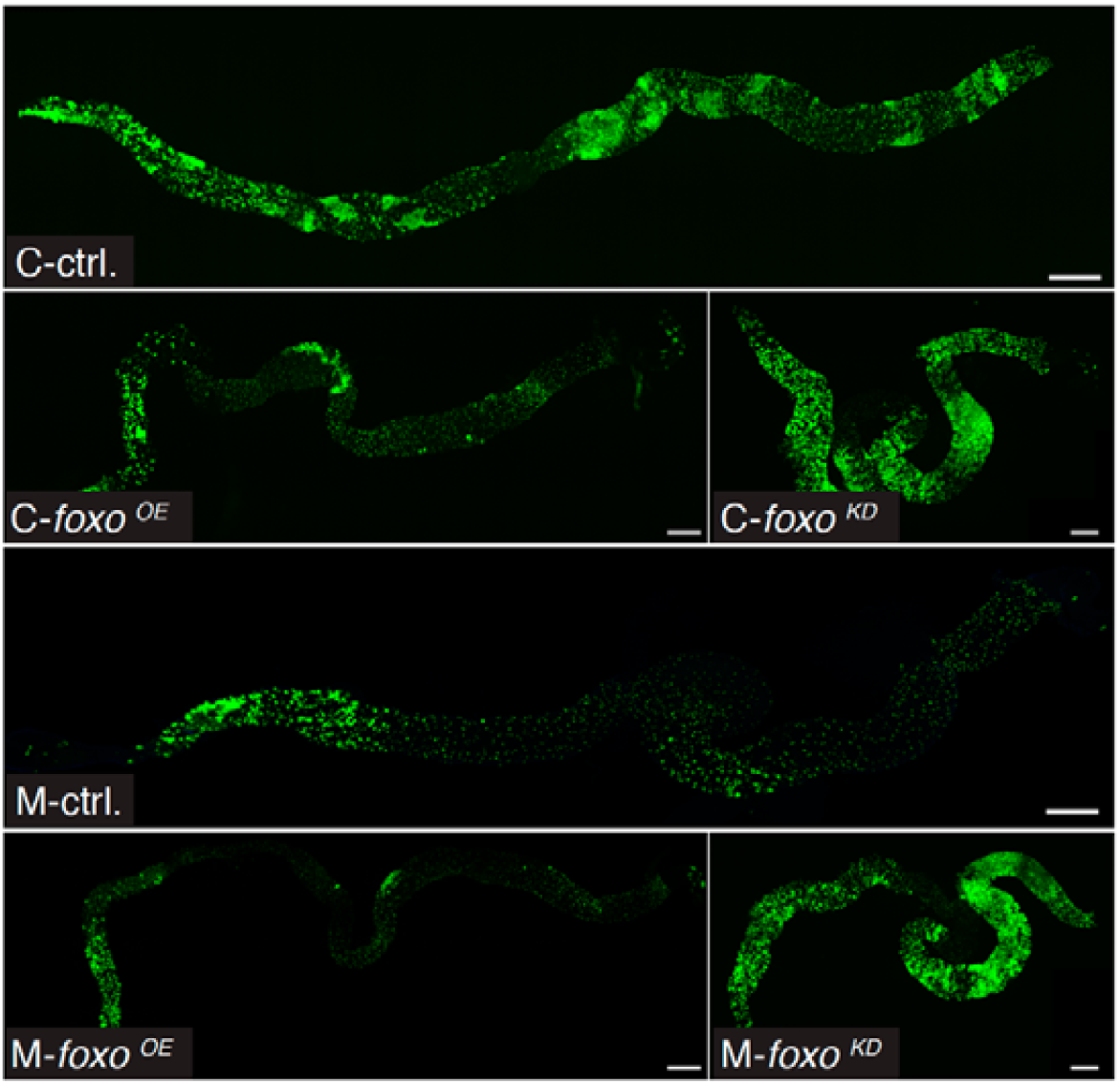
FOXO is involved in ISC differentiation defect. *foxo* overexpression in ISCs (*esg*^*ts*^*F/O>mitoXhoI, foxo*^*OE*^ flies) reduced the GFP^+^ midgut area in the C group midguts, while *foxo* knockdown increased the GFP^+^ area in the M group (*esg*^*ts*^*F/O>mitoXhoI, foxo*^*KD*^ flies). Assessed 10 days after heat shock. Scale bars: 200 µm.

